# Plasma pTau-217 Correlates with Brain Atrophy, Cognition, and CSF Biomarkers in a Cognitively Healthy Community Cohort

**DOI:** 10.1101/2025.04.30.651583

**Authors:** Ming Ann Sim, Ella Rowsthorn, William T. O’Brien, Mujun Sun, Lachlan Cribb, Katherine Franks, Stuart J. McDonald, Stephanie Yiallourou, Marina Cavuoto, Ian H. Harding, Trevor T-J Chong, Meng Law, Terence J. O’Brien, Lucy Vivash, Matthew P. Pase

## Abstract

Plasma biomarkers are promising for detecting Alzheimer’s disease (AD) pathology, but their role in cognitively healthy individuals remains unclear. Plasma pTau-217 has high diagnostic accuracy for clinical and prodromal AD, yet its relevance in preclinical stages is underexplored. We examined if plasma biomarkers of AD, neurodegeneration, and neuroinflammation were associated with cognition, brain structure, and their cerebrospinal fluid (CSF) counterparts in dementia-free older adults. We studied community-based, dementia-free older adults from the Brain and Cognitive Health (BACH) cohort. Neuropsychological testing assessed global cognition (MMSE), memory (Logical Memory II), visual processing (Hooper Visual Organization Test), processing speed (Trail Making Test-A), and reasoning (Similarities). Paired plasma and CSF biomarkers (pTau-217, pTau-181, GFAP, NfL, Aβ42/40) were measured using SIMOA. MRI-derived cortical thickness was used as a neurodegeneration marker. Multivariable linear regression assessed associations between log_10_-transformed plasma biomarker levels, cognition (adjusted for age, sex, education, hypertension, hyperlipidemia, diabetes), and cortical thickness (adjusted for age, sex, education, and intracranial volume). Pearson’s correlations and Bland-Altman plots evaluated plasma-CSF agreement. There were 147 dementia-free participants (mean age±SD: 66.7±7.7 years; 56 % women). Higher plasma pTau-217 levels associated with lower global cognition scores (β -0.80, 95% C.I. -1.56, -0.03, p=0.041) and abstract reasoning (β -0.86, 95% confidence interval [C.I.] -1.62, -0.09, p=0.028). Greater plasma pTau-217 also associated with lower global cortical thickness (βeta [β] -0.21, 95% confidence interval [C.I]. -0.37, -0.06, per log unit change; p=0.006). No associations were found between the plasma biomarkers and processing speed or visual processing (p>0.05 for all). Among 47 participants with paired plasma-CSF biomarkers, plasma pTau-217 showed the strongest correlation with its CSF counterpart (R=0.76, p<0.0001), outperforming pTau-181 (R=0.61, p<0.0001), GFAP (R=0.66, p<0.0001), NfL (R=0.56, p<0.0001), and Aβ42/40 (R=0.53, p=0.0001). In conclusion, plasma pTau-217 levels were associated with both cognition and cortical thickness in dementia-free older adults. All plasma biomarkers correlated significantly with their CSF counterpart. These findings reinforce the utility of plasma biomarkers, particularly pTau-217, as indicators of neurodegenerative processes, even in asymptomatic individuals.

## INTRODUCTION

Biofluid biomarkers of Alzheimer’s disease (AD), neurodegeneration, and neuroinflammation provide valuable insight into neurodegenerative processes. They may support early diagnosis of AD pathology and help stratify individuals at risk of dementia ^1^. Whereas cerebrospinal fluid (CSF) biomarkers provide a direct window into these processes^2^, their invasive measurement precludes widespread scalability ^3,4^. Plasma biomarkers offer a minimally invasive alternative, although their ability to detect preclinical neurodegenerative changes in cognitively healthy individuals remains underexplored ^5^.

Plasma pTau-217 has emerged as a highly sensitive biomarker of AD pathology^6^. However, its relevance in cognitively normal individuals remains unclear. Indeed, in addition to pTau-217, emerging biomarkers within the National Institute on Aging and the Alzheimer’s Association (NIA-AA) framework for the diagnosis and characterization of AD also include biomarkers reflective of diverse neuropathology such as neurodegeneration (Neurofilament Light, NfL), neuroinflammation (Glial Fibrillary Acidic Protein (GFAP)), pTau-181 as well as Aβ42/40 (amyloid)^5^. Therefore, evaluating plasma pTau-217 and these related biomarkers in a dementia-free community-cohort may reveal early neurodegenerative changes and help identify individuals at risk before clinical symptoms emerge^7-10^.

To this end, we aimed to investigate if plasma biomarkers of AD (pTau-217, pTau-181, Aβ42/40), neurodegeneration (NfL), and neuroinflammation (GFAP) were associated cross-sectionally with cognitive performance, cortical thickness (as a proxy of early structural neurodegenerative processes)^11,12^, and their CSF counterparts in a dementia-free, community-based cohort. Investigating these associations will allow for a better understanding of these biomarkers in the context of a community setting, providing useful insight for refining early detection and prevention efforts for Alzheimer’s disease.

## MATERIALS AND METHODS

### Study cohort design

An overview of the study design is presented in **Figure 1**. The Brain and Cognitive Health (BACH) cohort is an ongoing prospective observational cohort study of Australian dementia-free community-based participants living in Australia, conducted at Monash University and the Alfred Hospital. Participants were recruited from August 2022 to November 2023. Participants were recruited via media and community advertisements. Written informed consent was obtained from all participants in accordance with the Declaration of Helsinki, and the study was approved by the Alfred Health Ethics Committee (Project ID 532/21). The inclusion criteria were adults aged 55-80 years, with at least 6 years of formal education, able to provide informed consent, and with sufficient language, visual and auditory acuity to complete cognitive testing. The exclusion criteria included any history of Schizophrenia, bipolar disorder, or schizoaffective disorder; current (or recent, defined as being within 2 years) major depression or within the last two years; alcohol abuse; diagnosis of dementia; significant neurological illnesses including multiple sclerosis, Parkinson’s’ disease or a disabling stroke (minor stroke was not an exclusion). One informant for each participant was also enrolled in the study, providing information on participant activities of daily living and cognitive functioning. All cognitive, functional, medical, demographic, and informant information was reviewed by a study dementia review committee comprising the study PI, a clinical neuropsychologist, and cognitive neurologist. This study reports on cross-sectional data from the first 147 participants.

**Figure 1.**
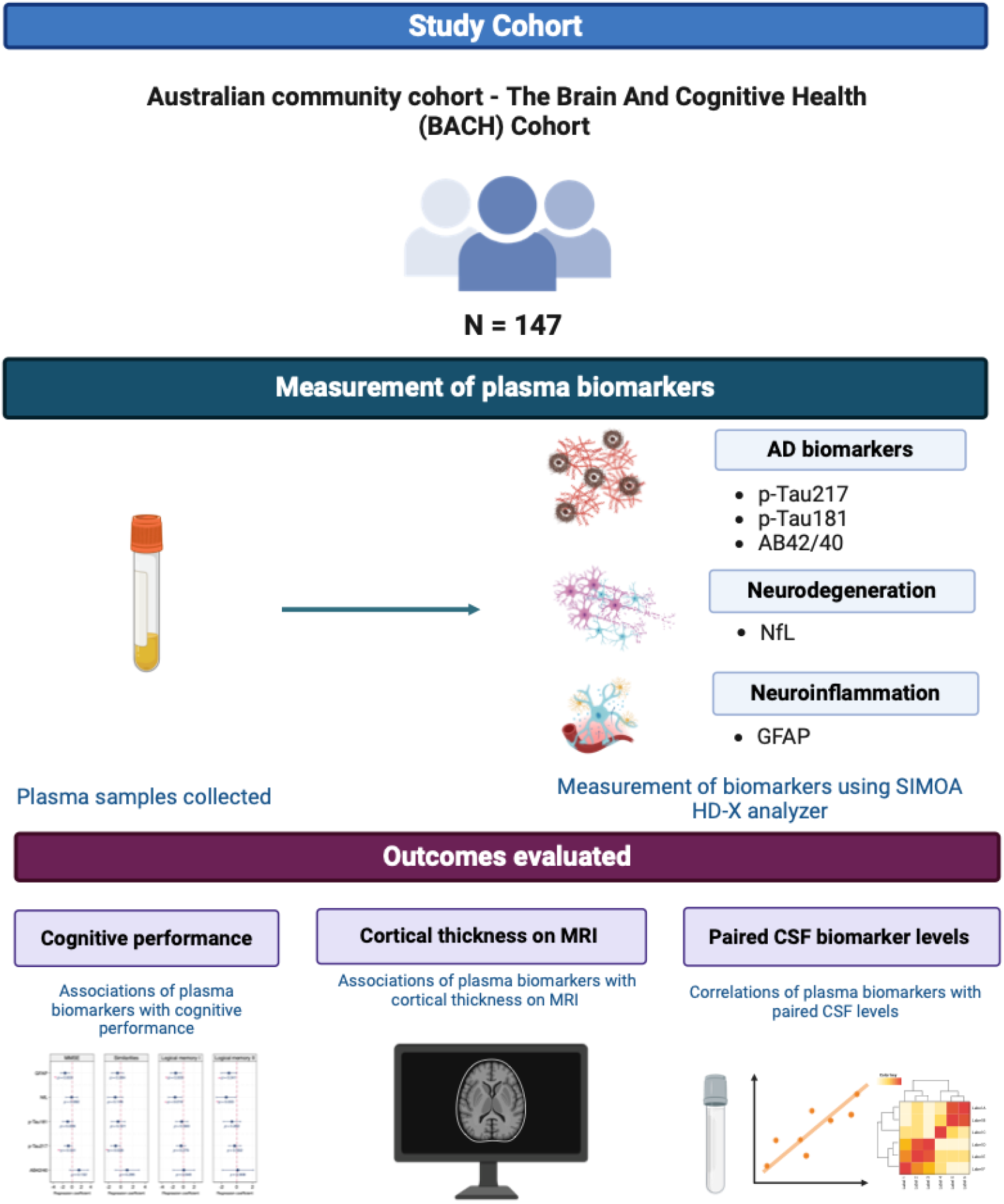
Overview of study.

### Clinical variables

Participant clinical and demographic data regarding age, sex, education level and medical history were obtained via interview questions administered in a standardized fashion by trained research assistants.

### Neuropsychological assessments

All participants underwent comprehensive neuropsychological testing, including two phone assessments and an in-person battery lasting approximately one hour. For the purpose of this study, we focused on the Mini-Mental State Examination (MMSE) as a measure of global cognition, Logical Memory II subtest (Wechsler Memory Scale; WMS IV) for episodic memory, the Similarities subtest (Weschler Adult Intelligence Scale IV) for abstract reasoning, the Hooper Visual Organization Task (HVOT) for visuospatial ability, and the Trail Making Test Part-A (TMT-A) for processing speed^13-16^. Performance on each test was standardized by creating *Z*- scores. All neuropsychological tests were administered at the Alfred Hospital, Melbourne, apart from Similarities, which was administered via telephone. Higher scores reflect better performance, except for TMT-A where higher scores reflect slower completion times.

### Neuroimaging assessments

Brain magnetic resonance imaging (MRI) scans were performed with the 3T Siemens Prisma Scanner at the Alfred Centre, Melbourne, including a T1-weighted scan: 3D magnetisation-prepared rapid gradient-echo (MPRAGE) sequence with TR 2400ms, TE 2.24ms, flip angle 8°, FOV 240 x 256 x 154mm, 192 sagittal slices of 0.8mm thickness, resolution 0.8 x 0.8 x 0.8mm. T1w images were then pre-processed using FreeSurfer, a freely available and automated software tool used to quantify volumetric brain neuroimaging measures^17^. Global cortical thickness was measured using Free-Surfer and used as a brain volumetric index of early neurodegeneration^12^. Estimated total intracranial volume (eTIV) was also estimated using FreeSurfer, to control for variability in head size during statistical analysis.

### Assessment of biofluid biomarkers

Fasting blood samples were collected on the morning of in-person cognitive and MRI assessments. Venous samples were collected within EDTA tubes with added prostaglandin E1, with plasma samples aliquoted, and stored at -80 degrees prior to use. Paired CSF samples were obtained for a subset of participants who provided informed consent for the lumbar puncture procedure. The intervertebral space below the level of the spinal cord (L3/4 and below) was punctured using a pencil point atraumatic needle, with CSF fluid collected in a standardized fashion, in polypropylene tubes. CSF samples were then centrifuged at 2000g at 4 degrees for 10 minutes, following which, the supernatant pipetted into a separate polypropylene tube, and gently inverted to avoid gradient effects. CSF samples were then aliquoted into screw cap polypropylene tubes and stored at -80 degrees prior to use.

Quantitative immunoassays of plasma (and where available, paired CSF) levels of GFAP, NfL, pTau-181, Aβ40/42, and pTau-217 were measured using the SIMOA HD-X® analyser (Quanterix, Boston, Massachusetts, USA) in the Department of Neuroscience, Central Clinical School, Monash University. Biofluid levels of GFAP, NfL and Aβ40/42 were quantified using the SIMOA Human Neurology 4-Plex E (N4PE) assay, while p-Tau-181 was quantified using the SIMOA pTau-181 Advantage kit. pTau-217 levels were quantified using the SIMOA ALZpath pTau-217 Advantage Plus assays, in accordance with the manufacturer’s instructions. For the assessment of CSF pTau-217 levels, an additional 10-fold dilution was performed as per correspondence with the kit manufacturer. All paired CSF and plasma samples were run within the same plate to minimize inter-plate variation across participants. A single sample was run across all plates as an interpolate control. All samples measured above the manufacturer’s analytical lower limit of quantification (LLOQ). The LLOQ were 0.40 pg/mL for NfL, 2.89 pg/mL for GFAP, 0.378 pg/mL for Aβ42, 1.02 pg/mL for Aβ40, 0.00326pg/mL for pTau-217 and 2.00pg/mL for pTau-181.

### Statistical analysis

Statistical analysis was performed using R version 4.3.1 (R Core Team. 2023. R Foundation for Statistical Computing, Vienna, Austria) and STATA (StataCorp. 2023. Release 18. College Station, TX: Statacorp LLC). All tests were considered statistically significant if p<0.05. Patient characteristics were expressed as percentages for categorical variables and mean ± standard deviation (SD) for continuous variables.

Plasma biomarker levels were log-10 transformed prior to analysis due to skewness. Associations of log_10_-transformed plasma biomarker levels with individual cognitive test Z-scores were evaluated using multivariable linear regression adjusting for age, sex, education level (expressed as above or below tertiary education level), hypertension, hyperlipidemia, and diabetes. Associations of log-10 transformed plasma biomarker levels with MRI measures of cortical thickness were evaluated using linear multivariate regression. The model adjusted for age, sex, education level, and total intracranial volume.

Pearson’s correlations were employed to study correlations between paired CSF-plasma biomarker levels. Correlation coefficients and accompanying p-values were presented alongside scatterplots. Bland-Altman plots were employed to evaluate the agreement between paired samples, with mean differences, and 95% limits of agreement visualized.

Finally, bivariate correlations between AD biomarkers, and biomarkers of neurodegeneration and neuroinflammation were studied. Correlation coefficients and p-values adjusted for false discovery rate using the Benjamini and Hochberg method were visualized as heatmaps^18^.

## RESULTS

### Participant demographics

A total of 147 dementia-free participants were recruited and included in this analysis as part of the first data freeze. Cohort demographics are presented in **Table 1**. The mean age was 66.7±7.7 years, mean MMSE was 28.5±1.6, 93% of participants were Caucasian, and 44% were male.

**Table 1.** Participant characteristics of the BACH study cohort.

| **Characteristic** | **N=147**<br>**Mean ± SD / N (%)** | **N= 47**<br>**CSF biomarkers** |
| --- | --- | --- |
| **Age (years)** | 66.6±7.7 | 66.14±5.80 |
| **Education level, n (%)** |  |  |
| **Above tertiary level** | 92 (62.6) | 28 (59.6) |
| **Below tertiary level** | 55 (37.4) | 19 (40.4) |
| **MMSE , n correct** | 28.5±1.6 | 28.78±1.14 |
| **Similarities, n correct** | 27.2 (4.4) | 27.6 (3.8) |
| **HVOT, n correct** | 24.0 (3.3) | 24.0 (3.5) |
| **TMT-A, seconds** | 29.0 (8.9) | 27.1 (8.1) |
| **LM-II, n correct** | 20.0 (6.5) | 21.1 (5.7) |
| **Sex (women), N (%)** | 97 (66.0) | 26 (55.3) |
| **Race (Caucasian), N (%)** | 136 (93.0) | 45 (95.7) |
| **Diabetes, N (%)** | 9 (6.1) | 3 (6.4) |
| **Hypertension, N (%)** | 46 (31.3) | 12 (25.5) |
| **Hyperlipidemia, N (%)** | 52 (35.4) | 15 (31.9) |

Abbreviations: MMSE – Mini Mental State Examination. HVOT – Hooper Visual Organization Task. TMT-A – Trail Making Test Part-A. LM-II – Logical Memory II.

Mean fluid biomarker levels are presented in **Table 2**. Plasma biomarkers were available for all 147 participants, and MRI data was available for 145 participants. Paired CSF samples were available for a subset of 47 participants (mean age 66.14±5.80 years, mean MMSE 28.78±1.14). A flowchart of participant recruitment is presented in **Figure 2**.

**Table 2.**
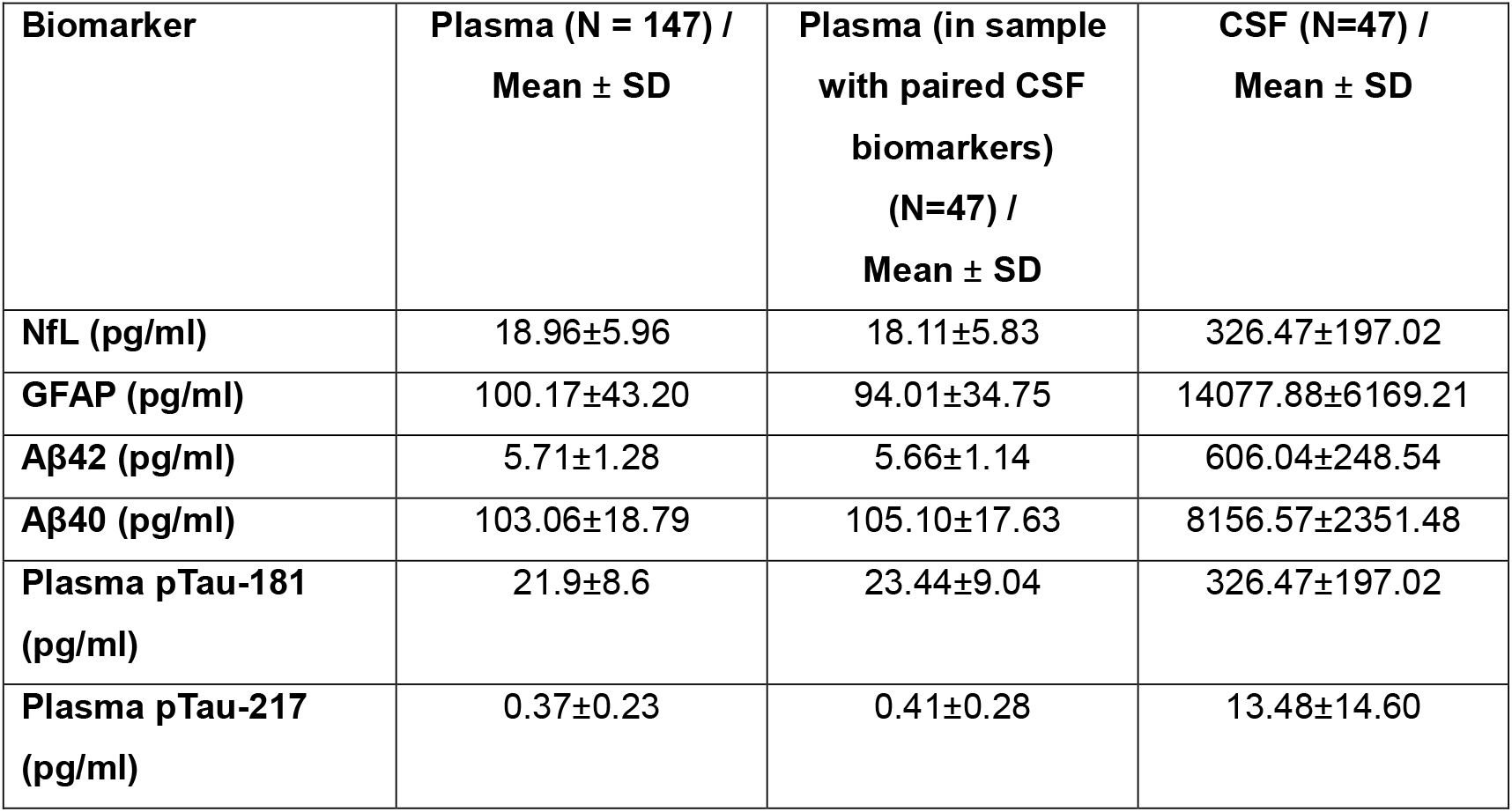
Mean plasma and CSF biomarker levels.

| **Biomarker** | **Plasma (N = 147) /**<br>**Mean ± SD** | **Plasma (in sample**<br>**with paired CSF**<br>**biomarkers)**<br>**(N=47) /**<br>**Mean ± SD** | **CSF (N=47) /**<br>**Mean ± SD** |
| --- | --- | --- | --- |
| **NfL (pg/ml)** | 18.96±5.96 | 18.11±5.83 | 326.47±197.02 |
| **GFAP (pg/ml)** | 100.17±43.20 | 94.01±34.75 | 14077.88±6169.21 |
| **Aβ42 (pg/ml)** | 5.71±1.28 | 5.66±1.14 | 606.04±248.54 |
| **Aβ40 (pg/ml)** | 103.06±18.79 | 105.10±17.63 | 8156.57±2351.48 |
| **Plasma pTau-181**<br>**(pg/ml)** | 21.9±8.6 | 23.44±9.04 | 326.47±197.02 |
| **Plasma pTau-217**<br>**(pg/ml)** | 0.37±0.23 | 0.41±0.28 | 13.48±14.60 |

**Figure 2.**
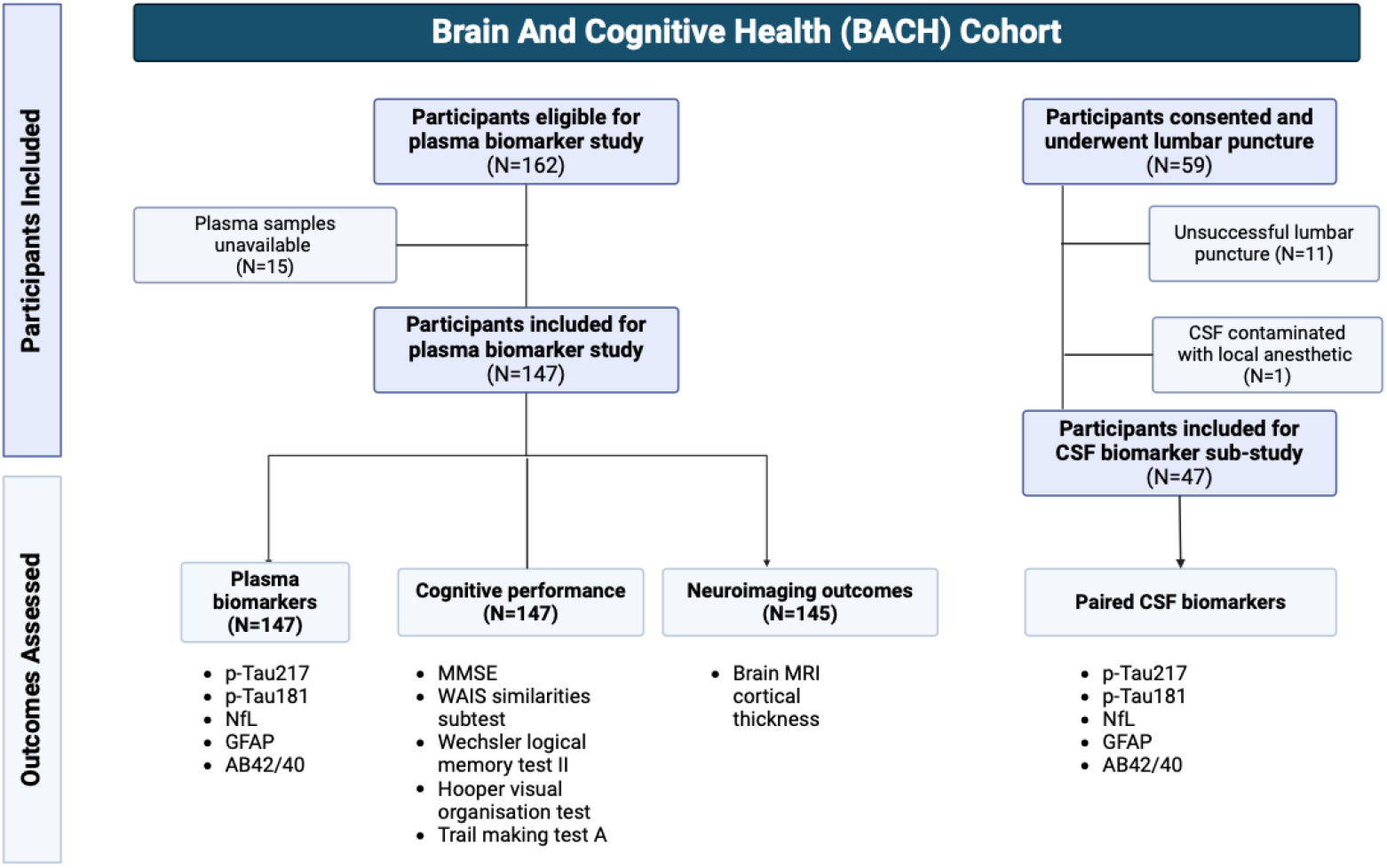
Flowchart of participant recruitment.

### Associations of plasma biomarkers with cognitive performance

The associations between the plasma biomarkers and cognitive tests are presented in **Figure 3A, and Table 3**, per log unit increase in each biomarker level. Higher levels of pTau-217 were associated significantly with poorer performance in global cognition (β -0.80, 95% C.I. -1.56, - 0.03, p=0.041) and the Similarities (abstract reasoning) subtest (β -0.86, 95% C.I. -1.62, -0.09, p=0.028). Meanwhile, higher levels of plasma GFAP were associated with poorer performance in global cognition (β -1.48, 95% C.I. -2.44, -0.52, p=0.003), and Logical Memory II tests (β - 1.05, 95% C.I. -2.05, -0.04, p=0.041). Higher plasma NfL levels were also associated with poorer performance on Logical Memory II (β-1.38, 95% C.I. -2.65, -0.12, p=0.033). Levels of GFAP, NfL, and pTau-217 were not significantly associated with performance on the HVOT (visuospatial ability) or TMT-A (processing speed) tests. Plasma Aβ42/40 and pTau-181 were not significantly associated with any cognitive tests.

**Table 3.** Associations of plasma biomarkers with cortical thickness on MRI, and cognitive performance (on MMSE, and tests of memory, visual processing, processing speed and verbal and abstract reasoning)

|  | **Beta** | **95% C.I.** | **p-value** |
| --- | --- | --- | --- |
| **Similarities\*** |  |  |  |
| pTau-217 | -0.86 | -1.62, -0.09 | **0.028** |
| pTau-181 | -0.45 | -1.55, 0.65 | 0.421 |
| GFAP | -0.54 | -1.53, 0.45 | 0.284 |
| NfL | -0.92 | -2.17, 0.33 | 0.148 |
| AB42/40 | 1.04 | -0.91, 2.98 | 0.295 |
| **HVOT\*** |  |  |  |
| pTau-217 | 0.12 | -0.68, 0.91 | 0.775 |
| pTau-181 | 0.24 | -0.89, 1.37 | 0.676 |
| GFAP | -0.11 | -1.13, 0.91 | 0.828 |
| NfL | -0.29 | -1.58, 0.99 | 0.653 |
| AB42/40 | -1.62 | -3.61, 0.36 | 0.108 |
| **TMT-A\*** |  |  |  |
| pTau-217 | 0.20 | -0.60, 1.00 | 0.629 |
| pTau-181 | 0.01 | -1.13, 1.15 | 0.988 |
| GFAP | 0.76 | -1.53, 0.45 | 0.284 |
| NfL | -0.20 | -1.50, 1.10 | 0.766 |
| AB42/40 | -0.65 | -2.67-1.37 | 0.524 |
| **Logical memory II\*** |  |  |  |
| pTau-217 | -0.27 | -1.06, 0.52 | 0.502 |
| pTau-181 | -0.64 | -1.77, 0.48 | 0.262 |
| GFAP | -1.05 | -2.05, -0.04 | **0.041** |
| NfL | -1.38 | -2.65, -0.12 | **0.033** |
| AB42/40 | 0.12 | -1.88, 2.11 | 0.908 |
| **MMSE\*** |  |  |  |
| pTau-217 | -0.80 | -1.56, -0.03 | **0.041** |
| pTau-181 | -0.92 | -2.02, 0.17 | 0.096 |
| GFAP | -1.48 | -2.44, -0.52 | **0.003** |
| NfL | -0.09 | -1.34, 1.17 | 0.892 |
| AB42/40 | 1.48 | -0.45, 3.42 | 0.132 |
| **Cortical thickness\*\*** |  |  |  |
| pTau-217 | -0.10 | -0.17, -0.03 | **0.008** |
| pTau-181 | 0.01 | -0.10, 0.12 | 0.899 |
| GFAP | -0.06 | -0.15, 0.04 | 0.224 |
| NfL | -0.05 | -0.17, 0.07 | 0.434 |
| AB42/40 | 0.11 | -0.09, 0.30 | 0.278 |

\*: Regression coefficients (beta) and 95% confidence intervals were obtained from linear regression adjusted for age, sex, education level, hypertension, hyperlipidemia and diabetes. Biomarker levels (expressed in pg/ml) were log-10 transformed prior to analysis. Cognitive test scores were transformed to z-scores prior to analysis. Higher cognitive test scores correspond to improved cognitive performance, except for TMT-A. Abbreviations: MMSE –Mini Mental State Examination. HVOT - Hooper Visual Organization Task. TMT-A – Trail Making Test Part-A.
\*\*.: Regression coefficients (beta) and 95% confidence intervals were obtained from linear regression adjusted for age, sex, education level, and intracranial volume. Biomarker levels (expressed in pg/ml) were log-10 transformed prior to analysis

**Figure 3.**
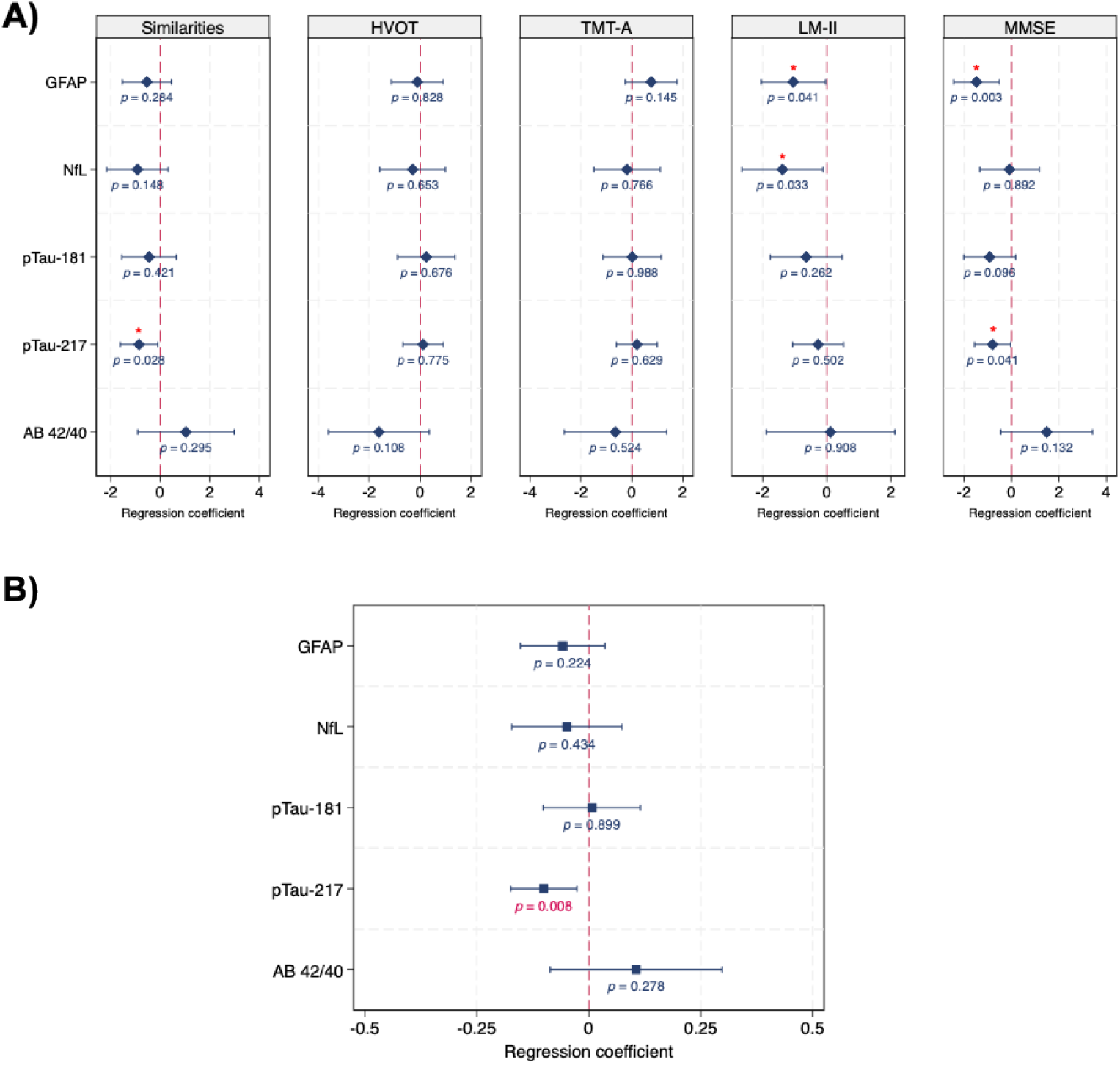
Associations of plasma biomarkers with A) cognitive performance and B) cortical thickness on brain MRI. “A) Regression (Beta) coefficients and 95% confidence intervals were derived from linear regression models adjusted for age, sex, education level, hypertension, hyperlipidemia, diabetes and log-10 transformed levels of plasma biomarkers (pTau-181, pTau-217, NFL, GFAP or AB42/40 respectively). B) Associations of plasma biomarkers with cortical thickness were derived from linear regression adjusted for age, sex, education, and intracranial volume. Figure 4 legend: A) correlation coefficients and p-values were derived from pairwise correlations of paired CSF and plasma biomarker levels. B) Bland Altman plots depict agreement between CSF and plasma biomarker levels, with mean differences and 95% limits of agreement visualized.”

### Associations of plasma biomarkers with brain cortical thickness

The associations between the plasma biomarkers and global cortical thickness are presented in **Figure 3B**, per log unit increase in each biomarker level. After adjustment for age, sex, education level and total intracranial volume, higher pTau-217 levels were associated significantly with lower cortical thickness (β -0.10, 95% C.I. -0.17, -0.03, p=0.008). No significant associations were found between the rest of the biomarkers (GFAP, pTau-181, NfL, Aβ42/40) with cortical thickness (**Table 3**).

### Agreement and correlations between paired CSF and plasma biomarker levels

Scatterplots showing the associations between paired plasma and CSF biomarker levels within the 47 participants are presented in **Figure 4A**. Plasma pTau-217 had the strongest correlations with its CSF counterparts (R:0.76, p<0.0001), followed by GFAP (R:0.66, p<0.0001), pTau-181 (R:0.61, p<0.0001), NfL (R:0.56, p<0.0001), and Aβ42/40 (R:0.53, p=0.0001).

**Figure 4.**
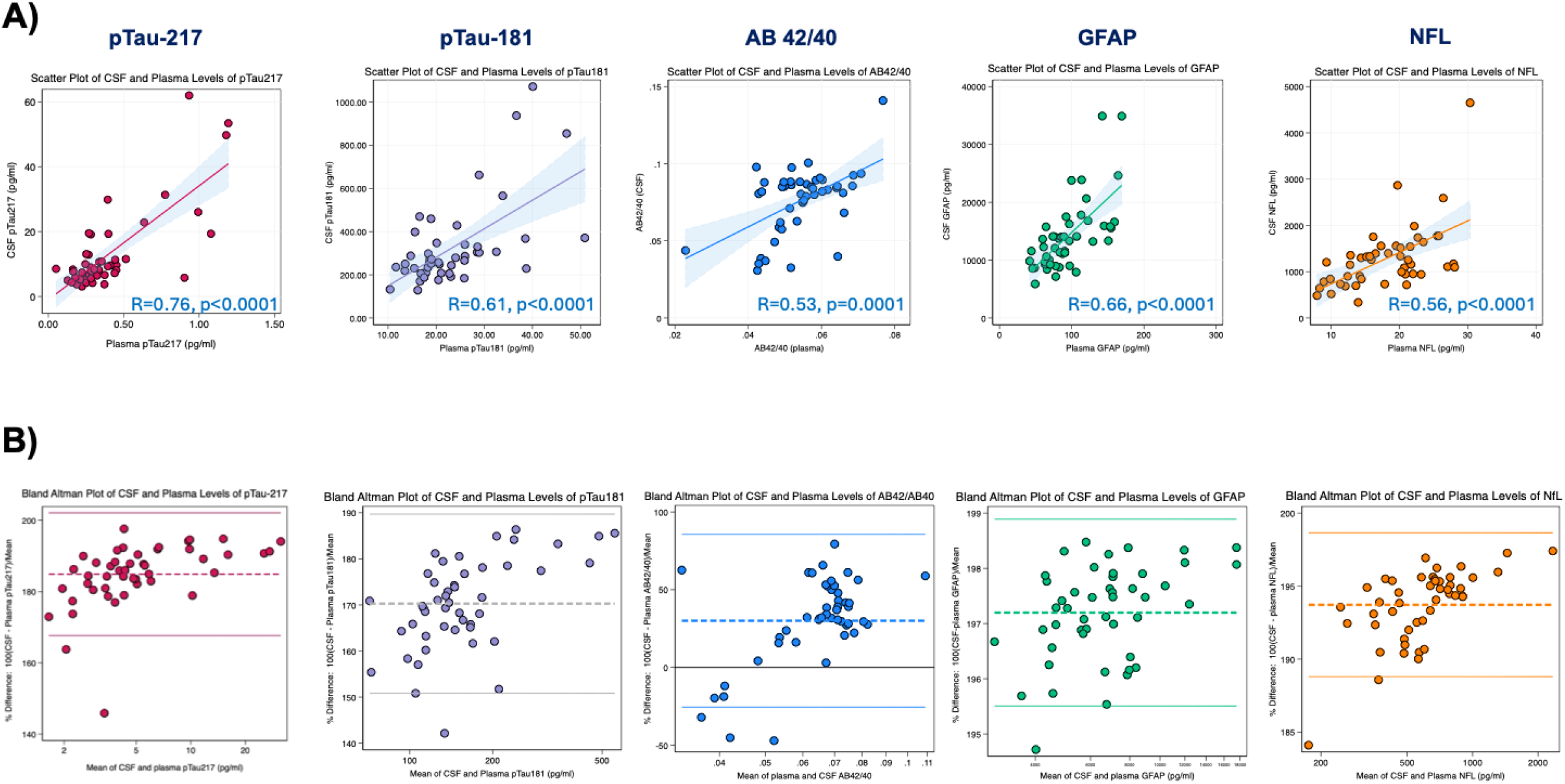
A) Scatterplots of paired CSF and plasma biomarker levels, B) Bland Altman plots of paired CSF and plasma biomarker levels (N=47)

Figure 4B presents Bland-Altman plots presenting the percentage mean difference and 95% limits of agreement between paired CSF and plasma biomarker values. Values falling outside the limits of agreement were mainly at the lower spectrum of measurement values. The highest agreement across both biofluids was observed for pTau-217, with values most closely centered around the mean. The greatest dispersion was observed for paired CSF and plasma Aβ42/40.

### Bivariate correlations between biomarkers of AD, neurodegeneration and neuroinflammation

Bivariate correlations between the fluid biomarkers are presented in **Figure 5**. Correlation coefficients and Benjamini-Hochberg adjusted p-values are presented in **Supplemental Tables S1 and S2**. CSF and plasma levels of GFAP and NfL significantly correlated with CSF and plasma levels of pTau isoforms. No significant correlations were found between Aβ42/40 and either GFAP or NfL.

**Figure 5.**
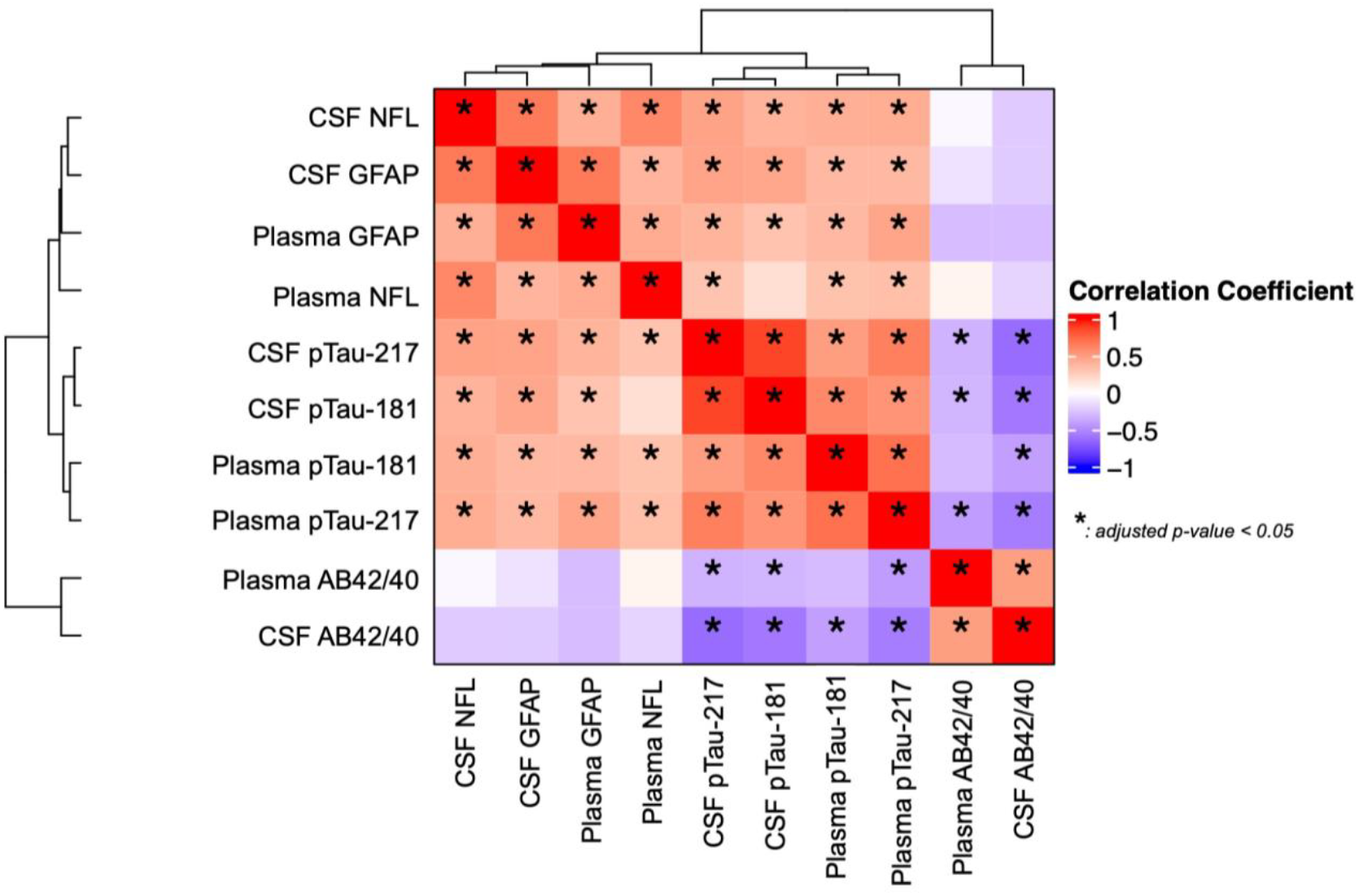
Bivariate correlations between biomarkers of AD, neurodegeneration and neuroinflammation (N=47) “Correlation coefficients were derived from bivariate correlations of biomarker levels. P-values were subject to false discovery rate correction using the Benjamini and Hochberg method.”

## DISCUSSION

In an Australian community-based cohort of dementia-free individuals, we found that higher levels of plasma pTau-217, NfL, and GFAP were associated with poorer cognitive performance (in particular, global cognition, abstract reasoning and memory domains) and were correlated with their CSF counterparts. Notably, higher plasma pTau-217 was also associated with lower cortical thickness. Although our results also demonstrated strong correlations between the plasma and CSF measures, pTau-217 demonstrated the strongest correlation with its CSF counterpart among all biomarkers examined. These findings reinforce the potential utility of plasma biomarkers, particularly pTau-217, as scalable and accessible indicators of neurodegenerative processes, even in asymptomatic individuals.

Our findings extend existing research, which has shown plasma pTau-217 to be a highly sensitive biomarker of AD pathology evaluated using PET and CSF biomarkers.^5,6^ We have broadened these findings to a dementia-free community-based cohort, showing that higher pTau-217 levels are already associated with reduced cognitive performance and cortical thinning before overt cognitive symptoms emerge. The stronger association of pTau-217 with a broader range of cognitive and neuroimaging markers, as compared to the other biomarkers, was expected and may be due to the extra sensitivity for early-stage amyloid pathology offered by pTau-217^6,19,20^ . Our observations are also consistent with previous reports demonstrating significant associations of plasma pTau-217 with brain atrophy in both AD and dementia-free patients^20,21^.

The associations between higher GFAP and NfL levels with poorer cognition further support their roles as disease-general markers of neuroinflammation and neurodegeneration, respectively ^22,23^. Our findings align with previous studies suggesting that NfL and GFAP may serve as early markers of cognitive dysfunction^9^. Indeed, these biomarkers have been associated with poorer cognitive performance during pre-clinical stages of dementia, with changes in their circulating levels occurring years prior to the onset of symptomatic cognitive disease^7-10^. Our results add to this body of evidence, demonstrating that even in a relatively healthy community-based cohort, elevations in these markers relate to cognitive function, particularly in global cognition and memory decline – cognitive areas which are broadly impacted by AD processes^24^.

Whereas pTau-217, GFAP, and NfL were significantly associated with cognitive performance in our cohort, pTau-181 and Aβ42/40 were not. Aβ and pTau biomarkers may be disease-specific and hence, vary in a time and pathology-dependent manner^25^. As such, pTau- 181 and Aβ42/40 may not capture the neuropathological burden in individuals without established AD who may have cognitive symptoms due to other processes, such as cerebrovascular pathology or Lewy body disease. Thus, these markers may be more specific to later-stage AD pathology. The poorer performance of immunoassay-based Aβ42/40 measurements as compared to mass spectrometry^26-28^ may have contributed to the weak associations observed between Aβ42/40 and the outcomes in the current study.

A major strength of our study was the head-to-head comparison of paired plasma and CSF biomarkers. Consistent with prior work, all plasma biomarkers studied showed moderate to strong correlations with their CSF counterparts.^29-34^ These findings provide support for the utility of plasma biomarkers as a proxy for central nervous system pathology, even in dementia-free populations.^30^ Of the blood biomarkers studied, pTau-217 demonstrated the strongest correlation and agreement with its CSF counterpart, reinforcing its utility as the most informative peripheral marker for detecting early neuropathological changes in the brain, relative to the other biomarkers examined. These findings are consistent with recent work demonstrating that CSF and plasma pTau-217 had the tightest agreement compared with that for pTau-231 and pTau-181^33^. Given its high diagnostic accuracy for tau and Aβ pathology, plasma pTau-217 may reduce the need for lumbar punctures when screening for AD risk and determining participant eligibility for disease-modifying therapy^5,35,36^.

Interestingly, we found that plasma pTau-217 more strongly correlated with CSF GFAP and NfL as compared with CSF Aβ42/40. This suggests that pTau-217 may more closely associate with astrocytic and neuronal dysfunction in the brain, as compared with Aβ42/40 in the central nervous system – a finding consistent with previous studies in MCI and AD cohorts^22,37^.

### Limitations

Our study has several limitations. First, the cross-sectional analysis precluded examination of how biomarker trajectories were associated with cognitive impairment. We are actively collecting longitudinal data to address this in the future. Secondly, whereas our sample includes a relatively healthy community-based cohort, validation of these findings in diverse populations with varying comorbidity burdens will help to enhance the generalizability of our results.^38^ Lastly, paired CSF and plasma levels of non ATN (Amyloid, Tau, Neurodegenerative) biomarkers relevant to dementia and co-existing pathophysiology such as inflammation should be studied. Indeed, it is recognized that while crosstalk exists across plasma and CSF, the inflammatory processes across the two entities may not be linear^38^. Nonetheless, our study has several strengths including the collection of paired CSF and plasma samples in a healthy cohort, facilitating a head-to-head comparison of several promising biomarkers. Our study, conducted within a community-based sample and with comprehensive neuropsychiatric assessment, highlights the role of these biomarkers in the pre-clinical setting, reinforcing their sensitivity to disease processes even in subclinical stages.

## Conclusions

In a dementia-free Australian community cohort, plasma pTau-217, NfL and GFAP were associated with cognitive performance and their paired CSF levels. Of the biomarkers studied, plasma pTau-217 displayed the strongest correlation with its paired CSF counterpart, and significantly associated with cortical thickness on MRI. Plasma biomarkers, particularly pTau- 217, are therefore, promising candidates for the detection of early neuropathology and central nervous system dysfunction even in older asymptomatic populations.

## Data availability

The data supporting the findings of this study are available from the corresponding authors, upon reasonable request.

## Acknowledgements

We thank the BACH study participants and their families for their dedication and commitment to the study.

